# Monotreme-specific conserved proteins derived from retroviral reverse transcriptase

**DOI:** 10.1101/2022.02.03.479050

**Authors:** Koichi Kitao, Takayuki Miyazawa, So Nakagawa

**Affiliations:** Laboratory of Virus-Host Coevolution, Institute for Frontier Life and Medical Sciences, Kyoto University, Sakyo-ku, Kyoto 606-8507, Japan; Department of Molecular Life Science, Tokai University School of Medicine, Isehara, Kanagawa 259-1193, Japan

## Abstract

Endogenous retroviruses (ERVs) have played an essential role in the evolution of mammals. Many ERV-derived genes are reported in the therians that are involved in placental development. However, the contribution of the ERV-derived genes in monotremes, which are oviparous mammals, remains to be uncovered. Here, we conducted a comprehensive search for possible ERV-derived genes in platypus and echidna genomes and identified three reverse transcriptase-like genes named *RTOM1*, *2*, and *3*. They were found to be clustered in the *GRIP2* intron. Phylogenetic analysis revealed that *RTOM1*, *2*, and *3* are strongly conserved between these species, and they were generated by tandem duplications before the divergence of platypus and echidna. The *RTOM* transcripts were specifically expressed in the testis, suggesting the physiological importance of *RTOM* genes. This is the first study reporting monotreme-specific *de novo* gene candidates derived from ERVs, which provides new insights into the unique evolution of monotremes.

## Introduction

Endogenous retroviruses (ERVs) are remnants of retroviral genomes found in the host genomes. ERVs are retroviruses that infect the host germline cells and are integrated into the host genome (Johnson 2019). Young ERVs retain their viral open reading frames (ORFs), but gradually lose their intact ORFs due to the accumulation of mutations. However, proteins expressed from ERVs sometimes evolve as functional genes in the host (Ueda et al., 2020). A typical example is the syncytin genes, ERV-derived fusogenic genes, which are expressed in the human placenta (Mi et al., 2000; Blond et al., 2000; Blaise et al., 2003) and are required for mouse placenta formation (Dupressoir et al., 2009; Dupressoir et al., 2011). Syncytin genes have been independently acquired from different ERVs in different mammalian lineages, which is a representative example of the convergent evolution (Imakawa et al., 2015). In addition, other ERV-derived genes that does not show fusogenic activity have also been found to be expressed in the placenta. For example, *HEMO* encoding a secreted envelope protein (Heidmann et al., 2017) as well as *gagV1* and *pre-gagV1* genes (Boso et al., 2021) are highly expressed in the human placenta. In contrast, it is unknown whether ERV-derived genes are co-opted in monotremes that are egg-laying mammals.

Here, we attempted to determine whether there are ERV-derived genes specific to monotremes. Comparative studies for the detection of ERV-derived genes have been conducted in mammalian genomes, including the platypus (Nakagawa and Takahashi 2016; Wang and Han 2020). However, for monotremes, only the genome sequence of one species, the platypus, was available (OANA5), the quality of which was limited (Warren et al., 2008). Recently, high-quality monotreme genomes of platypus (mOrnAna1.p.v1) and echidna (mTacAcu1.pri) were sequenced using long-read sequencing technology (Zhou et al., 2021). Taking advantage of these genome sequences, we conducted comparative analyses and detected three novel ERV-derived genes specific to the monotreme lineage.

## Results and Discussion

To comprehensively search for ERV genes in monotremes, we extracted ORFs from the genomes of platypus and echidna. The amino acid sequences obtained by the virtual translation of these ORFs were used as queries for the sequence search. We used the hidden Markov model (HMM) of the retroviral genes in the Gypsy Database 2.0 (GyDB) (Llorens et al., 2011) as the subject of the sequence search (Supplementary file 1—Table S1). We identified ORFs similar to *gag*,*pro*, *pol*, and *env* genes (Figure 1A). These ORFs are presumed to be a mixture of (1) ORFs that are evolutionarily conserved and (2) ORFs of young transposons that retain their ORFs. To exclude young ERV ORFs, we performed the clustering analysis based on the amino acid sequence identity. Since young ERVs are thought to be included in large clusters due to their mutual similarity, we removed sequences that belonged to large clusters consisting of more than 10 sequences. This step could also exclude evolutionarily conserved but highly duplicated genes such as SCAN domain-containing genes (Emerson and Thomas 2011), which is out of the scope of this study. Next, using the platypus ORFs as queries, and the echidna ORFs as the subjects, we conducted a sequence similarity search using BLASTp. We obtained ORF pairs with high amino acid similarity (Figure 1—figure supplement 1A; Supplementary file 1— Table S2). One of these ORFs was *ASPRV1* that is a known ERV-derived protease gene acquired in the common ancestor of mammals and is responsible for skin maintenance (Matsui et al., 2011). Next, we focused on the ORFs that are absent in human genome and identified three ORFs. They were located tandemly in the intron of the *GRIP2* gene in the opposite direction (Figure 1B). All three ORFs showed high similarity to the reverse transcriptase (RT) of spumaretrovirus in GyDB (Supplementary file 1 — Table S3). Therefore, we designated these genes as *RTOM* [RT-like ORF in Monotreme], and three genes were named as *RTOM1*, *RTOM2*, and *RTOM3* in order of their location from the 5’ direction (Figure 1B). To rule out the possibility that the *RTOM* genes were acquired before the divergence of humans and monotremes and were lost in humans, the *RTOM* coding sequences were searched using tBLASTx with in the genomes of six mammals, two birds, eight reptilians, and two amphibians (Supplementary file 1 — Table S4). Significant hits excluding the RT regions were not obtained other than in platypus and echidna. Therefore, the *RTOM* genes were thought to be acquired after the divergence of therians and monotremes.

**Figure 1.**
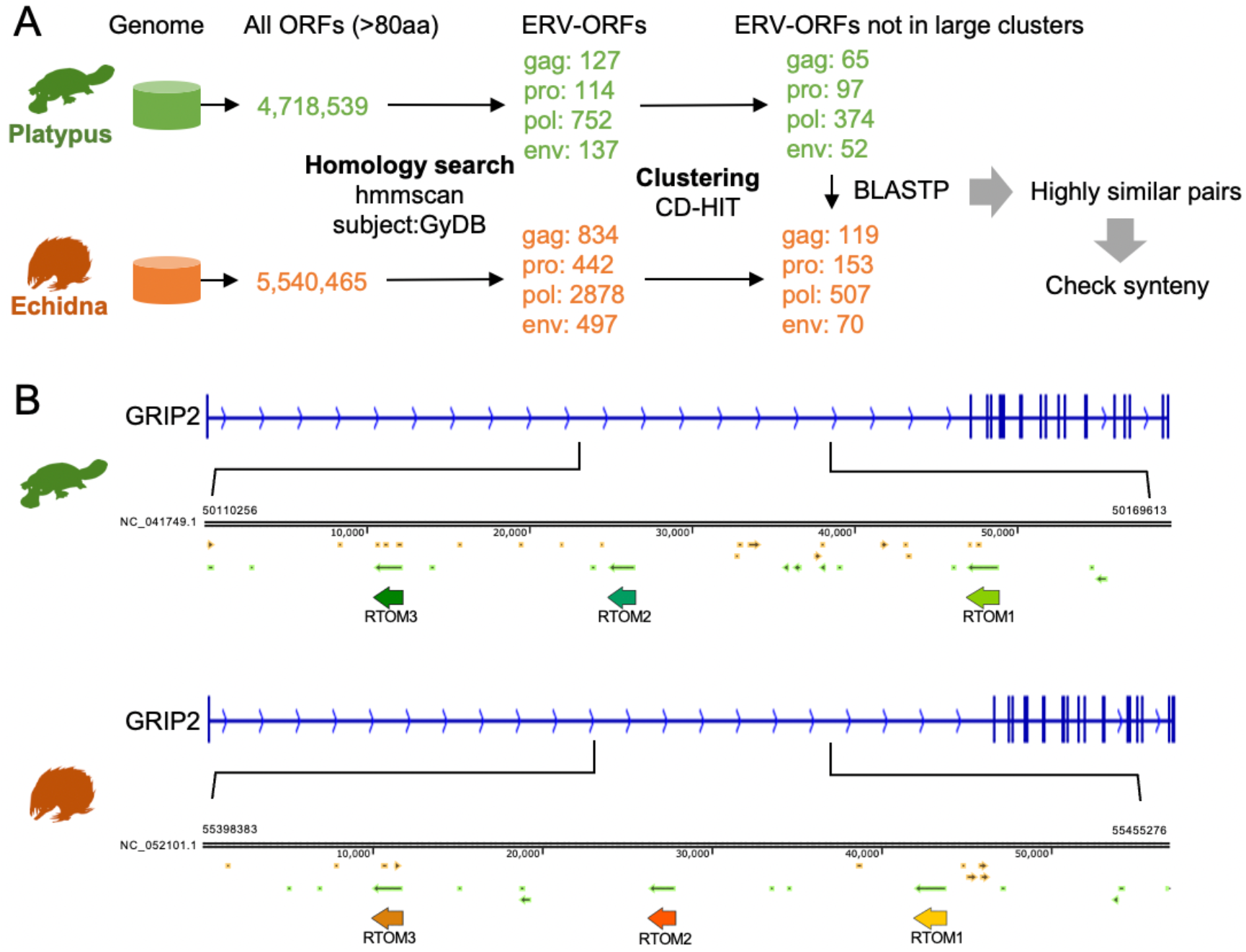
Identification of *RTOM1*, *2*, and *3*. (A) Schematic representation of the in silico screening for conserved ERV-derived genes in platypus and echidna. (B) Genomic context of *RTOM1*, *2*, and *3*.

We found that the *RTOM* genes in the platypus genome were computationally annotated in the RefSeq database (Figure 2A). *RTOM1*, *2*, and *3* genes of platypus contained two introns in the 5’ UTR, and the entire *RTOM* ORFs are expressed as mRNA excluding a second splicing variant of *RTOM3* that partially lost its ORF (Figure 2A). In echidna, *RTOM2* and *RTOM3* gene structures were annotated in the RefSeq transcripts; however, *RTOM1* was not annotated. By conducting transcriptome assemblies of RNA-seq data of echidna tissues (Supplementary file 1 — Table S5), we reconstructed the *RTOM1* transcript (Figure 2B; Supplementary file 2). As a result, all echidna *RTOM* transcripts have two introns in the 5’ UTR, which was similar to observations for platypus. According to the alignment of the six amino acid sequences of platypus and echidna RTOM proteins, RTOM2 lacks a region shared by RTOM1 and RTOM3, but the C-terminal region was conserved among the RTOM proteins without insertion or deletion (Figure 2C). To investigate the tissue-specific expression of *RTOM* genes, we analyzed the RNA-seq data of platypus and echidna (Supplementary file 1—Table S5). In platypus, *RTOM1*, *2*, and *3* were commonly highly expressed in the testis (Figure 2D). *GRIP2* was expressed not only in the testis but also in the brain, and its expression level was lower than that of the *RTOM* genes. This suggests that the *RTOM* expression was not a result of the *GRIP2* expression. We further investigated the mapped reads using Interactive Genome Viewer (Thorvaldsdóttir et al., 2013). It was found that *RTOM3* showed a splicing variant with an intron in the coding region, as shown in the RefSeq transcript (Figure 2—figure supplement 1). In echidna, we found that all *RTOM* transcripts were specifically expressed in the testis, similar to platypus. Expression of *GRIP2* in echidna testis was also relatively low, strengthening the idea that the *RTOM* expression is independent of *GRIP2* expression (Figure 2E). Given the higher expression level of *RTOM2* in both platypus and echidna, this gene may play a central role of the RTOM proteins. It is still possible that the relative expression levels of three genes may change according to tissues and developmental stages that were not examined in this study.

**Figure 2.**
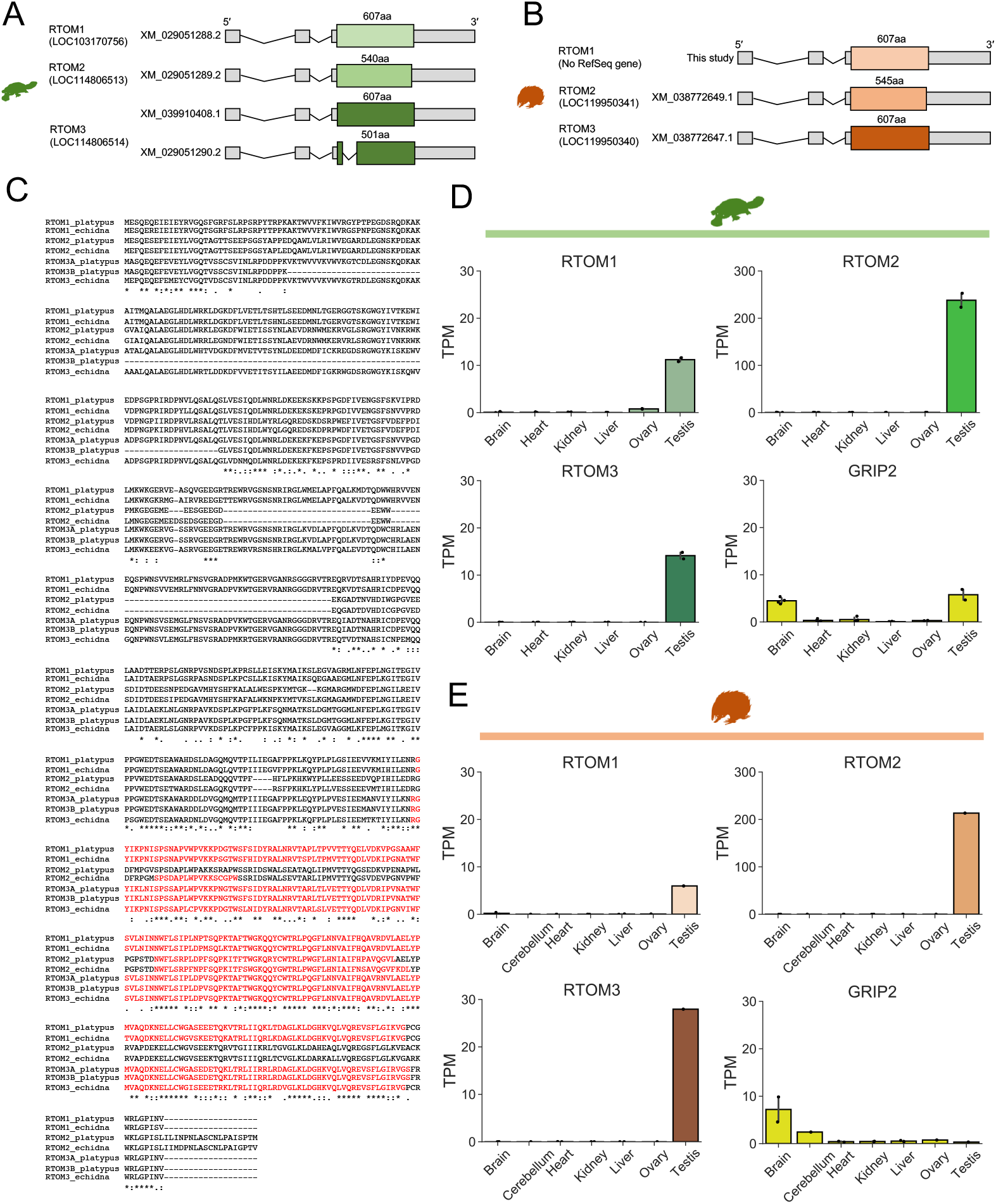
Expression of *RTOM1*, *2*, and *3*. (A) Schematic representation of the RefSeq transcripts of the *RTOM* genes in platypus. (B) Schematic representation of the reconstructed *RTOM1* transcript and RefSeq transcripts of the *RTOM2* and *3* genes in echidna. (C) Multiple alignment of the amino acid sequences of RTOM proteins. The amino acid sequence of echidna RTOM1 was obtained from the genomic ORF. “RTOM3A_plasypus” and “RTOM3B_platypus” are protein isoforms derived from “XM_039910408.1” and “XM_029051290.2,” respectively. The regions showing similarity to the HMM of spumaretrovirus RT domain in GyDB are indicated in red. (D and E) Tissue-specific expression of *RTOM* genes and *GRIP2* in (D) platypus and (E) echidna. Normalized expression levels are presented as transcript per million (TPM).

To obtain insights into the viral origin of the *RTOM* genes, we performed a BLASTp search of the amino acid sequence of platypus RTOM1 against the NCBI virus database. We found that retrovirus Pol proteins from various distinct lineages, namely gammaretrovirus, deltaretrovirus, epsilonretrovirus, and spumaretrovirus, are similar to the RTOM1 proteins (BLASTp: E-value < 1E-20). In all hits, the retroviral Pol proteins showed high similarity to the latter half of RTOM1 (about 370-607aa) (Figure 3A). Domain search against the Pfam database (Mistry et al., 2021) in the HMMER web service (Finn et al., 2011) revealed that the latter half of RTOM1 and RTOM3 contain RT domains (Figure 3—figure supplement 1). A phylogenetic tree was constructed from the RT regions of the RTOM proteins and the retroviral Pol proteins (Figure 3B). The RTOM proteins appear to be more related to class III retroviruses, including spumaviruses or spumavirus-related MuERV-L (Llorens et al., 2009). The tree topology of the RTOMs strengthens the validity of the inference that three *RTOM* genes were generated by tandem gene duplications before the divergence of platypus and echidna (Figure 3C). In the non-RT region of RTOM1 (approximately 1-369aa), no significant hits for retroviruses were obtained (Figure 3A). We performed a BLASTp search for all non-redundant proteins in the GenBank database for the non-RT region of RTOM1; however, no similar proteins were found except for RTOM2 and 3 (E-value < 0.05). Therefore, the non-RT region of the *RTOM* genes does not seem to be derived from ERV genes or conserved host genes. Considering the structural divergence of the non-RT region, such as deletion of RTOM2 and splicing variant of platypus RTOM3 (Figure 2C), the RT region is a core domain of the RTOM proteins, and the non-RT region may provide functional modifications specific to each RTOM protein.

**Figure 3.**
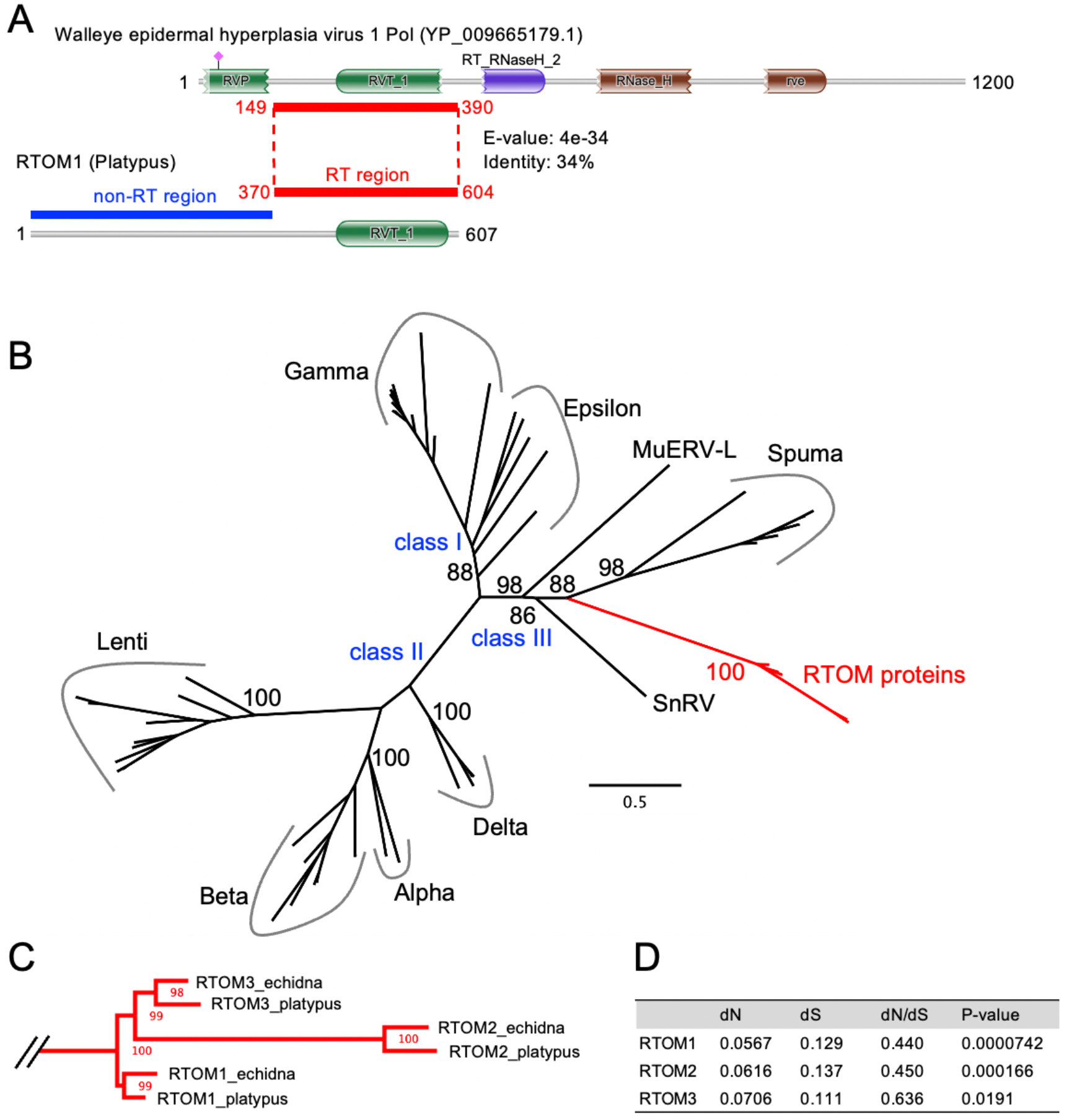
Evolution of *RTOM1*, *2*, and *3*. (A) Comparison between platypus RTOM1 and retroviral Pol protein. Walleye epidermal hyperplasia virus 1 is represented as an example. A region showing similarity to the Pol protein by BLASTp was designated as “RT region.” A region that did not show similarity to any retroviral genes was designated as “non-RT region.” (B) A phylogenetic tree constructed from the amino acid sequences of RT regions of the six RTOM proteins and the retroviral Pol proteins in GyDB. The multiple alignment is available in Supplementary file 3. Ultrafast-bootstrap values obtained from 1000 times replication are shown in major branches. (C) Detailed representation of the clade of the RTOM proteins. (D) The numbers of non-synonymous substitutions (dN) and synonymous substitutions (dS) per site estimated by Nei-Gojobori method (Nei and Gojobori 1986). Statistical significance of selection was estimated by the codon-based Z test of neutrality using MEGA-X (Kumar et al., 2018).

During the 187-million-years history after diverging from monotremes, therians have acquired many ERV genes and evolved their unique features, especially the placenta (Imakawa and Nakagawa 2017). Our work revealed that monotremes also domesticated ERV genes which emerged and was conserved more than 55 million years ago, the divergence time of platypus and echidna (Zhou et al., 2021). Calculation of nonsynonymous and synonymous nucleotide substitution frequencies of *RTOM1*, *2*, and *3* shows that they are under purifying selection, strongly suggesting their physiological importance (Figure 3D). One possibility of their functions is that they are restriction factors against ERVs and retrotransposons as well as exogenous retroviruses. For example, gag-derived *Fv1* (Best et al., 1996) and env-derived *Fv4* (Ikeda and Sugimura, 1989) inhibit retroviral infection in mice. It is possible that the RT domains in the *RTOM* genes may compete with retroviral infection and/or retrotransposition as antagonists. Another possibility is that RTOM proteins are involved in physiological functions unique to monotremes. To investigate that, it would be important to clarify which cells (viz. germ cells or somatic cells) in testis express the *RTOM* genes. Retroviral genes like *RTOM*, in which only RT domain is co-opted, have not been reported in other vertebrates to the best of our knowledge (Naville et al., 2016). Thus, the future functional elucidation of *RTOM1*, *2*, and *3* will provide us the new aspects of ERVs functioning in mammals.

## Materials and Methods

### Identification of conserved ERV genes

The platypus genome (mOrnAna1.p.v1, GCF_004115215.1) and the echidna genome (mTacAcu1.pri, GCF_015852505.1) were used for the ERV gene screening (please see fig. 1). The 240-nt ORF flanked by stop codons were retrieved using the getorf program in the European Molecular Biology Open Software Suite (Rice et al., 2000). For HMM-based sequence search, hmmscan was used (Expected threshold: 1E-5) in HMMER3 v3.2.2 (Eddy 2011). ORFs were clustered using CD-HIT v4.8.1 (Li and Godzik 2006) with 50% amino acid identity. The sequence search for platypus ORFs against echidna ORFs was conducted using BLASTp v2.10.0+ with an e-value < 1E-50 (Camacho et al., 2009). To detect the presence of homologous sequences beyond the monotreme lineage, deep homology searches were performed using tBLASTx v2.10.0+ with e-values <1E-5 on the genomes of six mammals, two birds, eight reptilians, and two amphibians (Supplementary file 1—Table S4)

To further examine the distribution of the bitscore of the BLASTp search, we extracted ORF pairs that showed high similarity (Figure 1—figure supplement 1A). Then, we examined the overlap with the RefSeq annotation and identified the genes to which the ORFs belonged. Finally, by extracting genes that had no homologs in the human genome, *RTOM1*, *2*, and *3* were detected. To validate this approach, we performed similar analyses on the genomes of human (GRCh38.p13) and marmoset (Callithrix_jacchus_cj1700_1.1) (Figure 1—figure supplement 1B and C). They diverged 43 million years ago (Perelman et al., 2011). We succeeded in identifying known ERV-derived genes such as *PEG10* (Ono et al., 2001), *RTL1/PEG11* (Charlier et al., 2001), *ASPRV1* (Matsui et al., 2011), *NYNLIN/CGIN1* (Marco and Marín 2009), *ERVV-1* and *2* (Kjeldbjerg et al., 2008), and *ERVMER34-1/HEMO* (Heidmann et al., 2017) (Supplementary file 1—Table S6). This suggests that our method is sensitive enough to identify ERV-ORFs conserved in platypus and echidna.

### Expression analysis

RNA-seq data of platypus (20 samples from 6 tissues) (Marin et al., 2017) and echidna (11 samples from 7 tissues) (Zhou et al., 2021) were used (Supplementary file 1—Table S5). Low-quality reads were trimmed and filtered using fastp v0.19.5 with default options (Chen et al., 2018). The filtered reads were mapped to each reference genome using HISAT2 v2.1.0 (Pertea et al., 2016). Based on the 11 RNA-seq sequencing data mapped on the echidna genome, we obtained the echidna *RTOM1* transcript by conducting transcriptome assembly using Stringtie2 v2.1.6 with “--merge” option (Kovaka et al., 2019). We added the coordinates of the echidna *RTOM1* transcript (Supplementary file 2) to the RefSeq gene coordinates. We then calculated the expression levels for 20 platypus and 11 echidna RNA-seq samples using the Stringtie2 program with default options (Kovaka et al., 2019).

### Phylogenetic analysis

Representative retroviral Pol amino acid sequences were retrieved from the GyDB collection (https://gydb.org/index.php/Alignment?alignment=POL_retroviridae_Biology_Direct_4_41_2009&format=txt) (Llorens et al., 2009). A multiple alignment was generated using MAFFT v7.487 (Katoh and Standley 2013), and poorly aligned regions were removed using trimAl v1.4.rev15 (Capella-Gutiérrez et al., 2009). A phylogenetic tree was constructed using IQ-TREE2 v2.0.8 (Minh et al., 2020) with 1000 replicates of ultrafast-bootstrap (Hoang et al., 2018). The tree was visualized using FigTree v1.4.4 (http://tree.bio.ed.ac.uk/software/figtree/).

## Acknowledgments

We would like to thank Editage for English language editing. This work was supported by Grant-in-Aid for JSPS fellows 20J22607 to K.K. and JSPS KAKENHI 20K06775 and 20H03150 to T.M. and S.N. The super-computing resource was partially supported by the NIG supercomputer at ROIS National Institute of Genetics.

---

**Figure 1—figure supplement 1.**
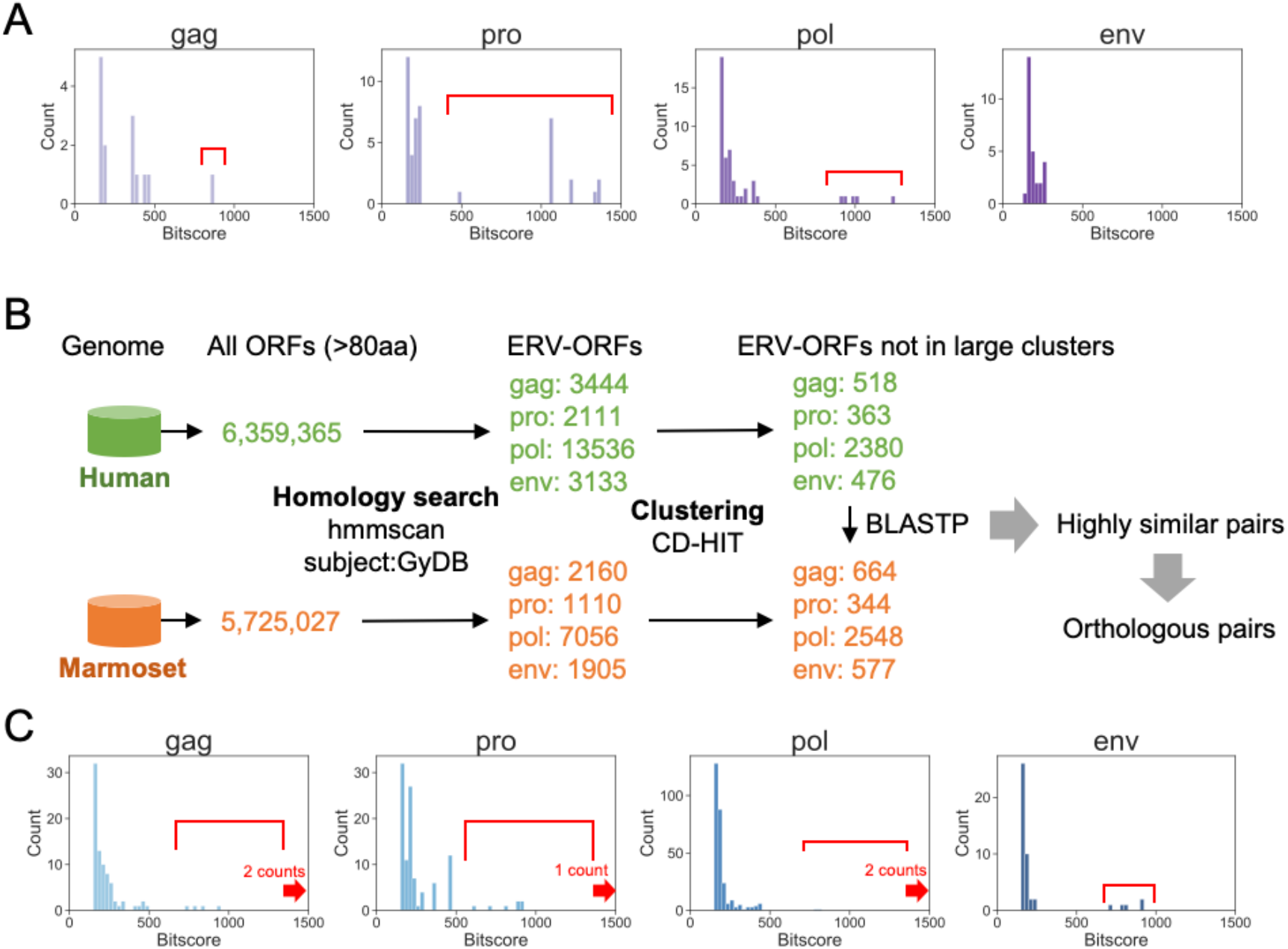
(A) Distribution of bitscore obtained by the BLASTp search for platypus ERV-ORFs against echidna ERV-ORFs. The ORF pairs with high bitscore indicated by red lines were used for further analysis (Supplementary file 1— Table S2). (B) Schematic representation of the control screening for conserved ERV-derived genes in human and marmoset. (C) Distribution of bitscore obtained by BLASTp for human ERV-ORFs against marmoset ERV-ORFs. The numbers of hits with bitscore higher than 1500 are indicated by red arrows. The ORF pairs with high bitscore indicated by red lines were used for further analysis (Supplementary file 1—Table S6).

**Figure 2—figure supplement 1.**
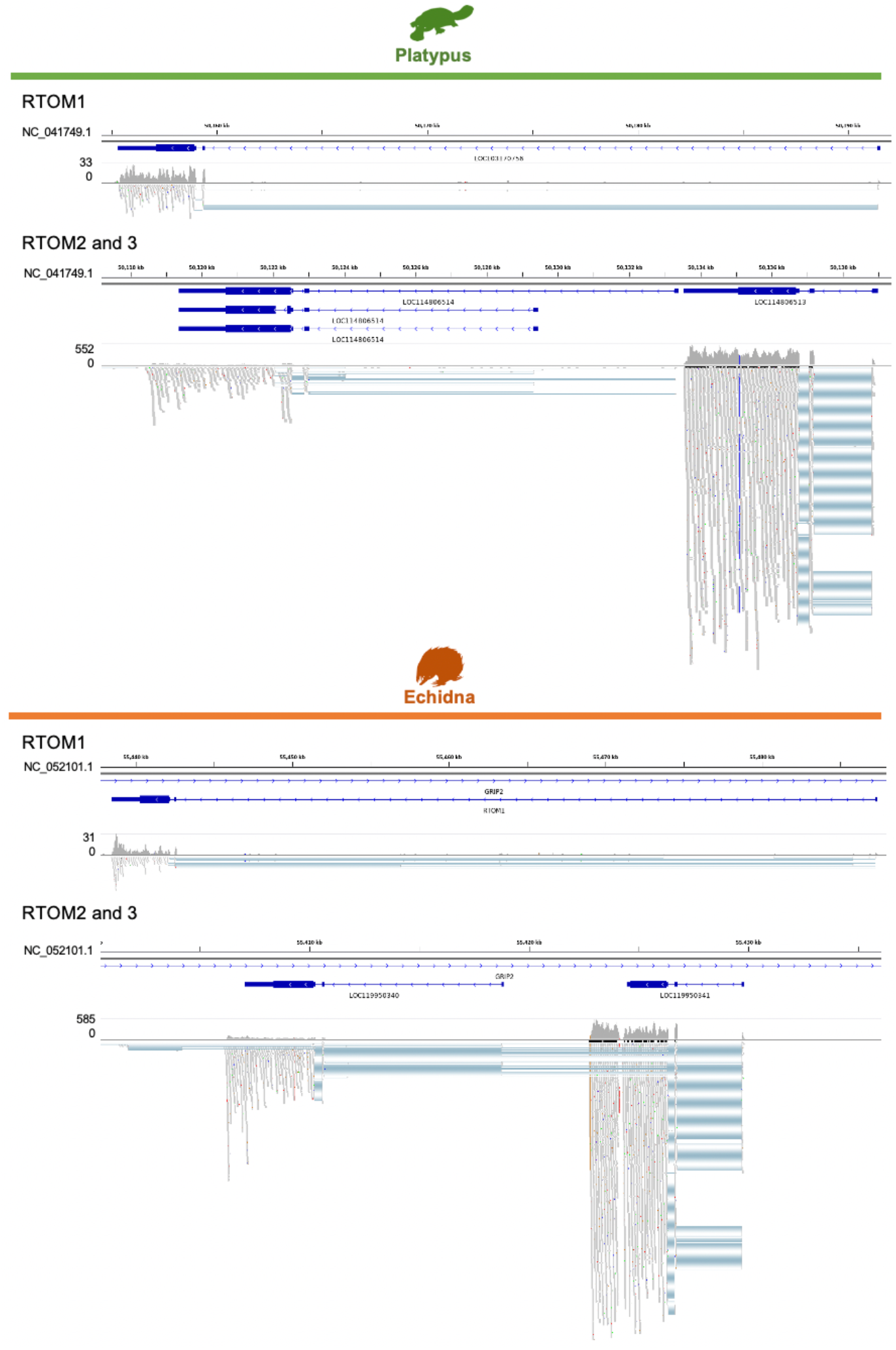
Screenshots of Interactive Genomic Viewer of RNA-seq reads on the *RTOM* genes. The transcript tracks in blue lines display the coordinates from the RefSeq GTF files. Thick blue lines indicate the coding sequences. Since there is no corresponding RefSeq transcript for echidna *RTOM1*, its gene coordinate was manually added from assembled transcripts in this study (Materials and Methods).

**Figure 3—figure supplement 1.**
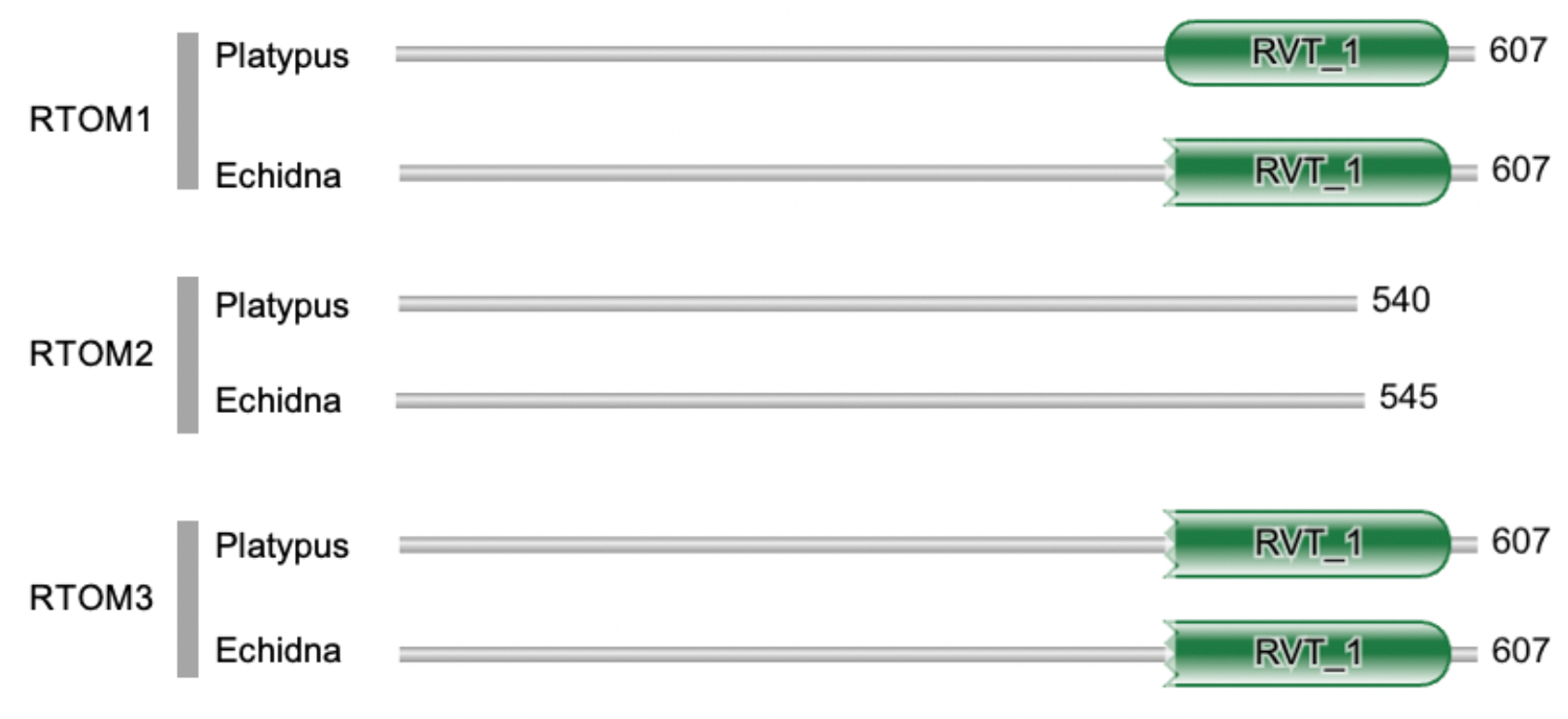
Protein domains in RTOM1, 2, and 3. The domain search was conducted using hmmscan in HMMER web server with default options (https://www.ebi.ac.uk/Tools/hmmer/search/hmmscan).

