## Supplementary data S1 for "Monotreme-specific conserved proteins derived from retroviral reverse transcriptase"

**Supplementary data S1.** Nucleotide sequence of the echidna *RTOM1* transcript

>RTOM1_echidna_exon1:NC_052101.1:55487268-55487386:-

ggaaaaggcagttagtttggttttggccttgctgtggcatgtgcactgcctaacggctcccttgagacccagccagaagacatctccacattgttaaagtaactctctcagcacttagg

>RTOM1_echidna_exon2:NC_052101.1:55442464-55442598:-

gacataacagccccttagaaactgagtcacagagtgctactgacagaaatttcttcagccactggccccagccaaaaagactaagaaaatagagcctcaaatccagatacaaaccaagggactgctcacattcag

>RTOM1_echidna_exon3:NC_052101.1:55438460-55442187:-

tttcagggtcctggctggcccagatctggtactgcaagcttcagaactgaggaccaagaccggatttcatcagccagcaATGGAATCTCAGGAGCGGGAGATTGAAATCGAGTACCGCGTGGGGCAAACTTCTGGCAGGTTTTCATTGCGGCCCTCAAGACCATATACTCCCCCAAAGGCTAAGACATGGGTGGTTTTTAAGATTTGGGTGAGGGGTTCCCCCAATCCAGAGGGTAACAGCAAGCAGGATAAGGCTAAAGCAATTACCATGCAGGCTTTGGCCGAGGGCCTACATGATTTGTGGCGTAAACTGGATGGCAAGGACTTTTTGGTAGAAACCCTAACGTCTCACAATTTGTCCGAAGAGGATATGAATTTGACAGGGGAGAGGGTTGGAACCAGCAAAGGGTGGGGATATATAGTAACTAAGGAGTGGATAGTGGACCCCAATGGTCCCAGGATTCGGGACCCATACCTACTGCAATCTGCCTTACAGTCTTTAGTAGAGAGCATTCAAGATTTGTGGAATAGACTGGATGAAAAGGAGAAATCTAAGGAGCCAAGACCTGGGGACATTGTAGTAGAAAATGGATCATTTTCCAATGTGGTCCCACGTGACCTAATGAAATGGAAAGGGAAGAGGATGGGAGCCATCAGAGTCAGGGAAGAAGGAGAAACTACAGAGTGGAGAGTGGGGAGCAATTCCAACAGGATTAGAGGCCTATGGATGGAACTAGCTCCCTTTCAAGCTTTAAAAGTGGACACTCAGGATTGGTGGCATAGAGTGGTTGAAAATGAGCAAAATCCTTGGAATTCTGTAGTAGAAATGAGATTATTTAACAATGTGGGTAGAGCTGACCCAGTGAAATGGACAGGAGAGAGGGTAAGAGCAAACAGAGGTGGGGGAGATAGAGTTACGAGGGAGCAGAAAGTGGACACCAGTGCTCACAGGATTTGTGACCCAGAGGTGCAACAACTTGCCATCGACACAGCAGAGAGACCAAGCCTAGGAAGTAGGCCAGCCTCAAATGATTCCCCTTTAAAGCCTTGCTCCCTGCTGAAAATTTCCAAATACATGGCTATTAAGTCGCAGGAGGGTGCGGCAGAGGGAATGTTGAATTTTGAGCCACTGAAGGGTATCACCGAGGGAATTGTCCCACCAGGGTGGGAAGACACCTCCGAAGCCTGGGCCCGTGATAACCTCGATGCTGGCCAGTTGCAGGTGACCCCCATCATTATAGAAGGGGTGTTTCCCCCTAAACTCAAACAGTACCCCCTTCCTTTGGGGAGTATTGAGGAAGTGGTTAAGATGATACATATCTTGGAGAATCGTGGCTACATAAAGCCAAATATTTCACCCTCCAATGCTCCAGTGTGGCCAGTAAAGAAGCCCAGTGGCACGTGGAGCTTCCATATTGATTATAGGGCTTTGAACAGAGTGACATCTCCATTGACTCCCATGGTAACCACCTATCAAGATTTAGTGGATAAGATCCCAGGGAATGCGACCTGGTTCTCAGTACTGAACATTAACAATTGGTTTTTGAGCATACCGCTCGACCCCATGAGCCAGCTTAAGACAGCTTTTACTTGGGGGAAGCAGCAATACTGCTGGACTAGGCTGCCTCAGGGATTTCTTAACAATGTGGCCATTTTTCATCAAGCCGTGCGGGACGTTTTAGCAGAGCTCTACCCCACGGTGGCCCAAGATAAGAATGAGCTCCTCTGCTGGGGGGTTTCTAAGGAGGAGACCCAAAAGGCAACCAGGCTCATTATCCAGAGATTGAAAGATGCGGGCCTCAAGCTTGATGGCCATAAAGTTCAGTTGGTTCAAAGAGAAGTGTCCTTTTTAGGAATCAAGGTTGGGCCTTGTGGATGGAGGCTGGGCCCTATCAGTGTTTAAcaaattgaagccttgcaatccacctggaatcggttcaaccctttaaaggactggtagcggttataagagttctatccccatgccgctgccttacaagccttttatagaaattggttaaaacacagaagtttcacaggatggattgttataacaataatagtgtcattagtaaagggtttactaataggagaaatggagtctagcagagaagagtctgatggaagcccttttaaaggccccggtgtagtggaccccaaaagtgagtcttccaggcatcctttatggagaggaatggggcagggagaggcctcccatccttattccccgatagaatggacagtttgagaatgccaaaggggattattgggaatatacagggcctttcagaagcttaaacacctgactggagagtgtgatgtgactgtgtagacaccccacgtcaccctataatccacctgggccaagctgctccagggaatcactctggtctgaagaggcatccccaaagctgatatggtgggctttggtactatccaatccccagatctggtaccataaattaaatgtgatggatcgaggggctctgggtattctcttgagatttagaatccattactgtggagcaggttaatatgggtccgccattctatattccgccattcctgtgtggtgaatggacttcaaggatcagggaatgatctatgctttgcccaagaatcacaattaagaacggggctttaaaaggggaaaccatccttttcccccaggaatactagacaactatcgccacggttgttttcatggagatactacaatcccaacatcctgccccatccatatagccccttttgtttgggcagacctctttctggaatttatcttccctctcgactgcctgacctctgcacttgagtctgcacctttgtacccccgaagtactcacactacccctgcagcacttttataaatgtccttatattggattccttcacctactgtaatttattttaatgtccacctccctggctagagtgtaaaccccttgagagcagagattgtgcctacttattctgttgtgccacccaacctttctcaaccaatgccaacttgcttgttctcaggttctttcagtaattggattgtttgctcatttgcctgttttgatatataacaaataggaggaggcgagtattactgttgtactagtttggatattgatggaaacacaacaagccactcccaagtcaagccatccagttgtgcaagtgaacaaggttgtaattcatcccttctggccatcgactttaaccaggatccttcctaatggttttcctctgataaataata
